# Predation of stocked salmon by riparian wildlife in semi-natural stream

**DOI:** 10.1101/127472

**Authors:** Kouta Miyamoto, Theodore E. Squires, Hitoshi Araki

## Abstract

Predation after release is one of the major concerns of hatchery fish conservation and propagation. However, the relationship among the size of hatchery fish, the predator species, and their behaviors in natural environments is largely unknown. To understand the relationship, we conducted predation experiments in outdoor tanks and a semi-natural stream with exposure to local predators. Two different ranges of fork lengths of masu salmon (*Oncorhynchus masou*) were examined as prey sizes. Camera trap data showed that grey herons (*Ardea cinerea*) were the primary predator animal in the system, and that most herons utilized shallow areas in the morning or evening. Increasing the density of stocked salmon brought in more grey herons. More importantly, predation by grey herons resulted in the survival rate of larger salmon being significantly lower than that of the smaller salmon. Our results suggest that it is important to understand local predators, adjust the optimum body size of hatchery fish at release, and choose the appropriate stocking site and time of day for maximizing the effectiveness of fish stocking.

## Introduction

Predation by riparian wildlife is widely recognized as a key factor influencing the survival of fishes in stream ecosystems (Kruuk, 1995; Draulans 1987; Roby *et al*. 2003; Steinmetz *et al*. 2003; Sahashi and Yoshiyama 2015). Salmonid fish populations reduced by riparian wildlife have been reported, not only for stocked populations, but also for wild populations (e.g. Osterback *et al*. 2013; Frechette *et al*. 2015). It is also noteworthy that stocked salmonids are often intensively preyed upon by birds (Wood, 1987; Martel and Dill, 1995; Harvey and Nakamoto, 2013) and mammals (Roberts and Garcia de Leaniz, 2011). Hatchery-reared salmonid fishes are routinely stocked in natural waterways as part of conservation efforts or stock enhancement (Brown and Laland 2003; Salvanes and Braithwaite 2006; Fraser 2008). Therefore, if we implement a management strategy that decreases local predation pressures, then effective conservation or stock enhancement methods should be improved.

Studies examining the predator-prey interaction between riparian animals and fish have been generally based on either investigating predation pressure of riparian animals without observing fish response (Draulans 1987; Post *et al*. 1998; Harvey and Nakamoto 2013) or investigating the mortality rate of fish without observing the behavior of predators (Penaluna *et al*. 2015). However, the predation effect is influenced by many factors: the condition of fish (Hostetter *et al*. 2012), fish migration (Roberts *et al*. 2009), the number of predators present, and the duration of their sojourn (Gawlik, 2002, Steinmetz *et al*. 2003). Thus, methodological improvements which decrease predation pressure on stocked fish require a profound understanding of both predator and prey behaviors. Additionally, as food size preference depends on predator species (Carss and Marquiss 1991, Sogard 1997) and the ability of fish to avoid predators depends on fish size (Dill and Fraser 1984), we need to consider the size effect of prey fish.

To evaluate the impacts of predation by riparian animals, it is necessary to evaluate the relationships among the predator species, their behaviors, and their preferred size of prey in a place that allows for free movement of all parties. However, the number of studies on this relationship is limited due to the difficulty in identifying species of predatory riparian animals and observing their predation behavior directly. These studies are further complicated due to complex behaviors exhibited by many predator species, such as individual movement for feeding and nesting needs (Collis *et al*. 2002), movement due to changes in prey densities (Kushlan, 1976; Gawlik, 2002), and movement resulting from human disturbance (Klein, 1993).

Recently, camera trappings that use fixed cameras, triggered by infrared sensors, to ‘trap’ images of passing animals, provide the opportunity to collect information including animals’ species, time of appearance in each day, appearance frequency, and duration of sojourn (Silveira *et al*. 2003; Wegge *et al*. 2004). Because camera trapping is a non-invasive technique, it causes minimal environmental disturbance (Henschel and Ray 2003; Silveira *et al*. 2003). In this study, we conducted a predation test using camera traps with outdoor experimental tanks and a semi-natural stream to investigate the relationships between the different predator species, their behavior, and the size of stocked salmon.

## Materials and Methods

### Test fish

The test fish used in the experiments were hatchery-reared masu salmon (*Oncorhynchus masou*) from the Shobu-shimizu River and the Jigoku River in Tochigi Prefecture, Japan (approximately 36°45'N, 139°27'E). Eggs were obtained from mature adults that had migrated into the Shobu-shimizu River and the Jigoku River, and were handled according to the standard hatchery procedures directed by the National Research Institute of Fisheries Science (NRIFS) facility at Nikko. Young of the year (YOY) (*n* = 305) and one year old (OYO) (*n* = 305) salmon were used in this study. The YOY and OYO salmon were graded from five rearing tanks (50 cm width × 120 cm length × 20 cm depth, the supply amount of water is set at 18 L/min) and a large tank (3 m width × 1.5 m length × 0.9 m depth, the supply amount of water is set at 150 L/min) respectively. The YOY fish used were 75–100 mm fish (75–100 mm FL-group, hereafter) and the OYO used were 135–160 mm fish (135–160 mm FL-group, hereafter). The body size range of the YOY salmon is bigger than that of general stocked YOY salmon in Japan (Sahashi *et al*. 2015), however the body size ranges are within those of the masu salmon that are stocked in the Shobu-shimizu River and the Jigoku River. In addition, the size range of YOY masu salmon are recommended as an effective body size for preventing predation by piscivorous fish in rivers around central Japan (Miyamoto and Araki 2017). Before the start of the study, fish were fed daily food rations (commercial trout pellets) equal to 1.5–2.0% of their estimated body weight.

### Tank experiment

We conducted predation tests in four outdoor tanks for three days from August to September, 2013. Four Fibre-Reinforced Plastic (FRP) circular tanks (120-cm diameter × 15-cm high) at NRIFS in Nikko were used in a forested and grassy area and each had a thin layer of natural gravel (2–7-cm gravel and about 13-cm cobble substrate) on the bottom (Fig. 1a). A 10-cm high and 8-cm roof fence that was installed on each tank, both to provide the fish with cover and prevented them from jumping out. A camera (Trophy Cam HD, Bushnell, Overland Park, KS, USA) was set in the south side of each tank to monitor the whole tank. The presence of cover and gravel substrate are set so that masu salmon can show near natural behavior, allowing hiding and escaping (Miyamoto 2016a), thus minimizing experimental stress. During the study days the stocked fish were fed daily food rations (commercial trout pellets) equal to around 2.0% of their body weight. Spring water, 10.2 ± 0.3°C (mean ± SD) was introduced into each tank at a rate of 6 L/min. Two different fork lengths (FL) of masu salmon were used as the size of prey fish, 75–100 mm FL-group (mean ± SD, 91.9 ± 7.0 mm) and 135–160 mm FL-group (145.8 ± 6.8 mm).

**Fig. 1.**
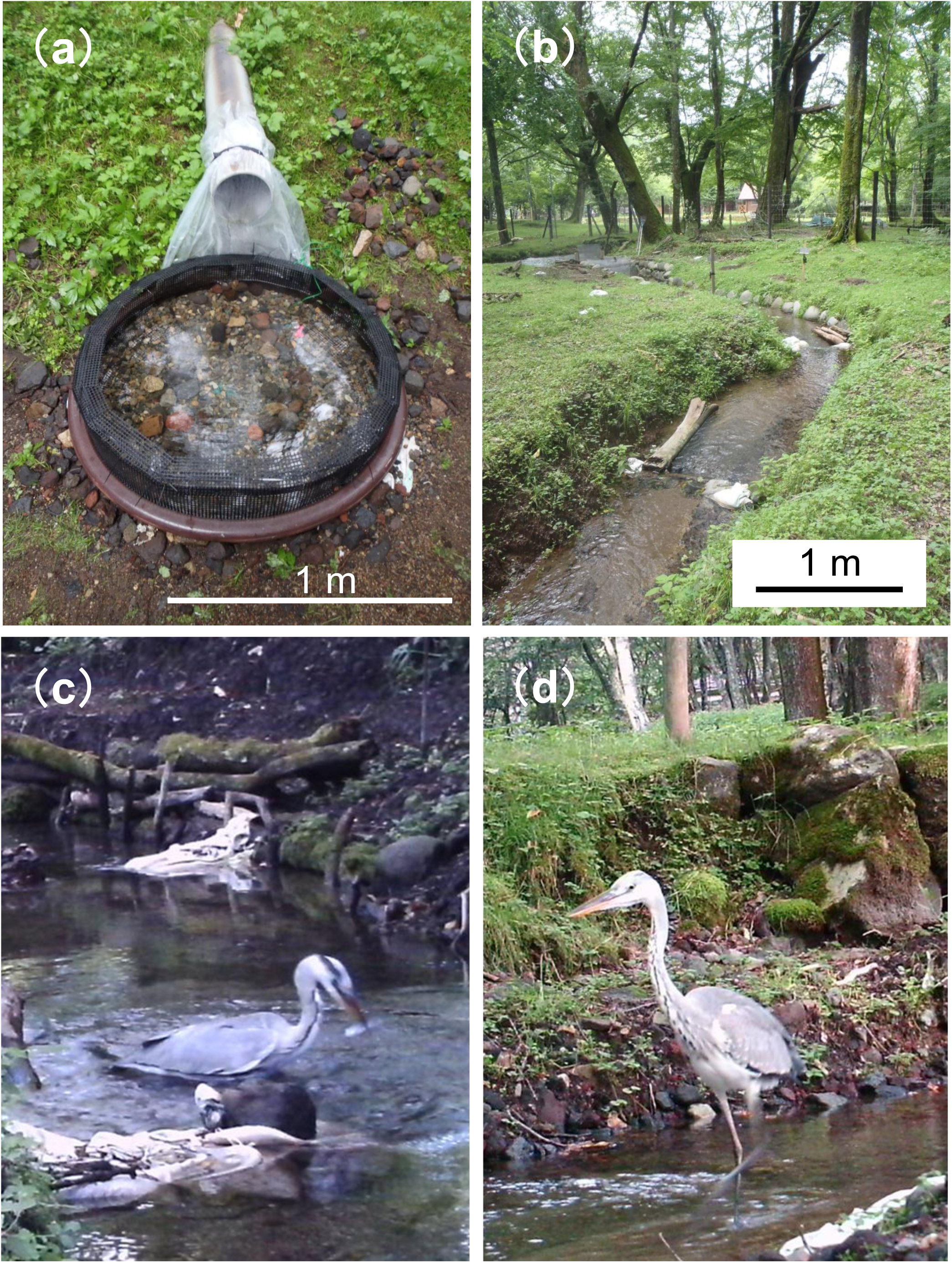
a and b: outdoor fibre-reinforced plastic circular tank (a) and semi-natural stream (b) at National Research Institute of Fisheries Science in Nikko. c and d: examples of photo of grey heron in the pool (c) and in the riffle (d) in the semi-natural stream.

To evaluate the size-selective predation risk, 30 fish (15 from each FL-group) were placed in each tank. The number of fish of each FL-group that survived was counted each day for three days. The survival rate of salmon was then compared between the two FL-groups. To identify the predator animals, the number of photos taken by camera trap (described in the section below) was counted.

Next, to investigate the relationship between the number of stocked salmon and the frequency in which predator animals appeared, three FRP circular tanks were used. The test fish were divided into 10, 5, and 0 individuals and stocked randomly in each of the three tanks. Three days later, the number of photos of potential predators taken by camera trap (described in the section below) was counted for each tank. This trial was replicated three times for each FL-group.

### Predation test in a semi-natural stream

The predation test was conducted for 20 days from August to September, 2013 using a semi-natural stream (mean ± standard deviation [SD]: 120 m long, 95.8 ± 4.3-cm wide; 2% gradient drain) in a forested and grassy area at NRIFS (Fig. 1b). The stream was constructed of stone, wood, and soil. Spring water, at 10.2 ± 0.3°C (mean ± SD) was drained at 18 L/s. The stream was forked to allow free passage into four 30 meter sections (Sec. 1-4 from upper to lower). Each section contained three pools and three riffles. The pools were about 80-cm long and about 60-cm wide with maximum depth of approximately 40-cm. Woody debris (c. 70-cm long and c. 20-cm wide) were placed in the sides of each pool. The riffles were 9–12-cm deep and contained a mixture of 2–7-cm gravel and about 13-cm cobble substrate. A 50-cm waterfall was built in the top of the stream with two metal gates (1-cm × 2-cm mesh) at the top to prevent fish from escaping. The water drain was separated by a metal mesh gate (1-cm × 2-cm mesh) with the bottom quarter of the gate covered with plastic mesh (8-mm × 8-mm mesh). A total of 400 fish (200 from each FL-group) were stocked in Sec. 2 after measuring their fork length and body weight. At the end of the study period, a backpack electrofishing unit (Smith-Root, Vancouver, WA, USA) was used to remove and count the fish that had survived. The electric shock was repeated until the fish count was zero twice in a row.

To investigate the distribution of salmon in the stream, the fish in each section were closed in by fish block nets (8-mm × 8-mm mesh) at the end of the experiment to prevent them from moving between the sections. When fish were caught by electrofishing, each fish was lightly anesthetized with 100 ppm 2-phenoxyethanol (Wako Pure Chemical Industries, Tokyo, Japan), its fork length and body weight were measured, and the section that the fish was caught in was recorded. To identify the predator animals, 3 cameras (Trophy Cam HD, Bushnell, Overland Park, KS, USA) were arranged in each section and placed at 10-m intervals along the riverside to monitor both sides of the stream, and the number of photos that were taken by camera traps (described in the section below) was counted for each section. In addition, to estimate the predation behavior of wild animals, the position of potential predators (at pool or riffle) in photos was recorded. Then, to investigate the position that predators utilized, the proportion of predators located around pools or riffles was calculated in each section, then the average proportions were compared.

### Camera trap

To assess predator encounters during day and night, potential predators were recoded using motion and infrared sensor camera traps in the tank experiments and the semi-natural stream experiment. Each camera was mounted on a wooden stake so that the camera was about 50 cm above the water’s surface. Cameras were triggered with a passive infrared motion sensor; the camera was set to wait 15 seconds after an initial trigger entered its sensor range before attempting to detect additional triggers. To identify predators and estimate the frequency of their visits to the study site, all the photos containing potential predators were checked by KM. For some ambiguous species identifications, additional checking was performed by a local wildlife expert Dr. T. Takeda from Nikko National Park. When more than one potential predator was captured in a photo, the species and their number was recorded. In addition, the number of photos that showed predators capturing or eating fish was counted, and the predator species was recorded.

### Statistical analyses

In the tank tests, Two-way ANOVA was used to determine the effects of study days and fish fork length on the survival of salmon. Additionally, a second set of ANOVA tests were used to determine how the number of salmon, and salmon size, effected the number of photos containing riparian predators in the stream test. One-way ANOVA was used to compare the average proportions of the predators located around pools and riffles in the stream. Count data that contain zeros was log (x + 0.5) transformed prior to analysis (Yamamura 1999), or, when proportions were tested, data were arcsin vx transformed. Tukey HSD test was used as a post-hoc test.

For the semi-natural stream experiment, Pearson's chi-square test was used to compare the proportion of the number of fish captured at each section. However, the omnibus chi-square value does not specify which combination of categories contributes to statistical significance; thus, the adjusted standardized residuals (ASR) was used for each value to determine discrepancies between the observed and expected value (Haberman, 1973). |ASR| > 1.96 and > 2.56 show *P* < 0.05 and *P* < 0.01, respectively. A *P* < 0.05 was considered significant in all statistical analyses. Data analyses were generated using IBM SPSS software (version 21.0 of IBM SPSS Statistics for Windows).

## Results

### Tank experiment

In the tank experiment, cameras captured grey heron (*Ardea cinerea*), Japanese marten (*Martes melampus*), raccoon dog (*Nyctereutes procyonoides*), and large-billed crow (*Corvus macrorhynchos*). The total number of photos containing potential predators was 186 (including three photos with ambiguous species, which were identified by the local wildlife expert). A total of 173 photos of grey heron were taken and there were six photos or fewer of each of the other animals (Japanese marten: three times, raccoon dog: four times, large-billed crow: six times). Twenty one photos showed grey herons capturing prey fish. Therefore, in tank experiments, grey herons were regarded as the main predator.

With regard to the size-selective predation risk, both the length of the study days and the fork length of the stocked salmon had significant effects on the fish survival (Period, *F*_2, 18_ = 10.56, *P* < 0.010; Salmon length, *F*_1, 18_ = 36.67, *P* < 0.001; Period × Salmon length, *F*_2, 18_ = 0.18, *P* = 0.833) (Table 1). The fish survival rate was significantly higher in the 75-100 mm FL-group than in the 135-160 mm FL-group during the study days (Tukey HSD test, all shows *P* < 0.050) (Fig. 2).

**Fig. 2.**
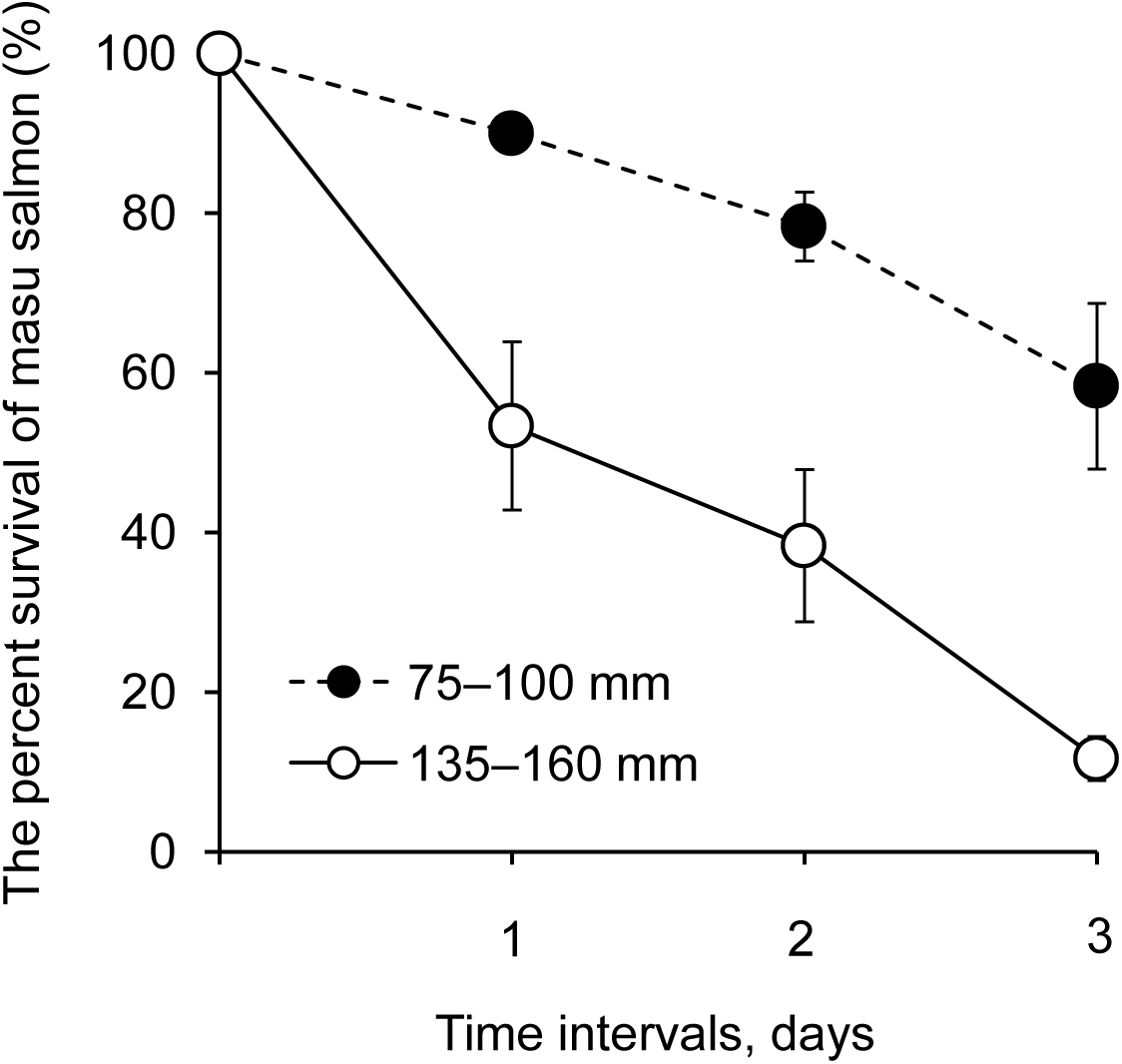
Survival rate of masu salmon (*Oncorhynchus masou*) with different body sizes groups in outdoor tanks. Circles and solid line: using 75–100 mm salmon; black circles and dashed line: using 135–160 mm salmon. Data are means ± standard error.

**Table 1.** Results from two-way ANOVA to determine the effect of study days, salmon length and their interaction on the survival of masu salmon (*Oncorhynchus masou*)

| Variable | *df* | *MS* | *F* | *p* |
| --- | --- | --- | --- | --- |
| Study days | 2 | 0.38 | 10.56 | **0.001** |
| Salmon length | 1 | 1.31 | 36.67 | **< 0.001** |
| Period × Salmon length | 2 | 0.01 | 0.18 | 0.833 |
| Error | 18 | 0.04 |  |  |

In the Two-way ANOVA test for frequency of predator appearance, only the number of fish in the tank had a significant effect on the number of photos containing grey herons (Number of fish, *F*_2, 12_ = 149.26, *P* < 0.001; Fork length, *F*_1, 12_ = 2.48, *P* = 0.141; Number of fish × Fork length, *F*_2, 12_ = 0.02, *P* = 0.981) (Table 2) and the number of photos containing grey herons increased significantly when the number of fish in the tanks was large (Tukey HSD test, all shows *P* < 0.001) (Fig. 3). At the end of the three-day experiment, there were no surviving salmon.

**Fig. 3.**
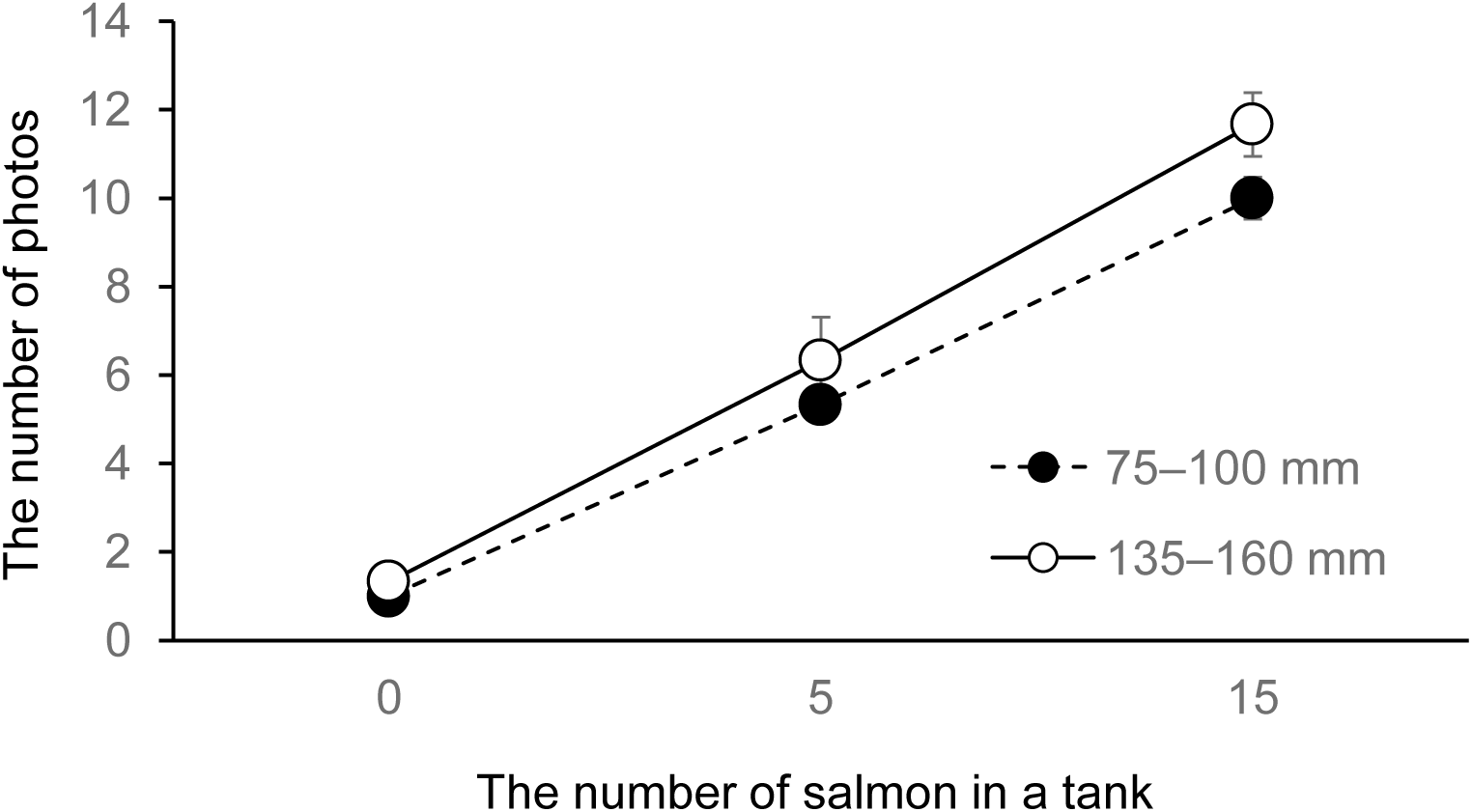
Relationship between the number of taken photos of grey heron (*Ardea cinerea*) and the number of masu salmon (*Oncorhynchus masou*) in outdoor tanks. Circles and solid line: using 75–100 mm salmon; black circles and dashed line: using 135–160 mm salmon. Data are means ± standard error.

**Table 2.** Results from two-way ANOVA to determine the effect of number of salmon, salmon length and their interaction on the number of photos containing grey herons (*Ardea cinerea*)

| Variable | *df* | *MS* | *F* | *p* |
| --- | --- | --- | --- | --- |
| Number of salmon | 2 | 1.11 | 149.26 | < **0.001** |
| Salmon length | 1 | 0.02 | 2.48 | 0.141 |
| Number of salmon × Salmon length | 2 | 0.00 | 0.02 | 0.981 |
| Error | 12 | 0.01 |  |  |

### Stream experiment

The overall survival of the fish in this experiment was 18.0% (*n* = 72). The survival of them was significantly higher for the 75–100 mm FL-group (33.0%, *n* = 66) than for the 135-160 mm FL-group (3.0%, *n* = 6) (Peason’s *χ*^2^ test, *χ*^2^ = 60.98, *df* = 1, *p* < 0.001).

There was no significant difference between the proportion of fish that survived in each section between both fish groups (Peason’s *χ*^2^ test, *χ*^2^ = 0.77, *df* = 3, *p* = 0.86). Comparing the actual proportion of surviving fish (75-100 and 135-160 mm) in each section to the expected surviving proportion (we hypothesized a uniform 1:1:1:1 ratio) showed there was a significant difference (Peason’s *χ*^2^ test, *χ*^2^ = 15.43, *df* = 3, *p* < 0.001). Accordingly, the number of fish surviving in Sec. 2 was significantly higher than the average number of fish capture in each section (ASR = 3.4, *p* < 0.010) (Fig. 4).

**Fig. 4.**
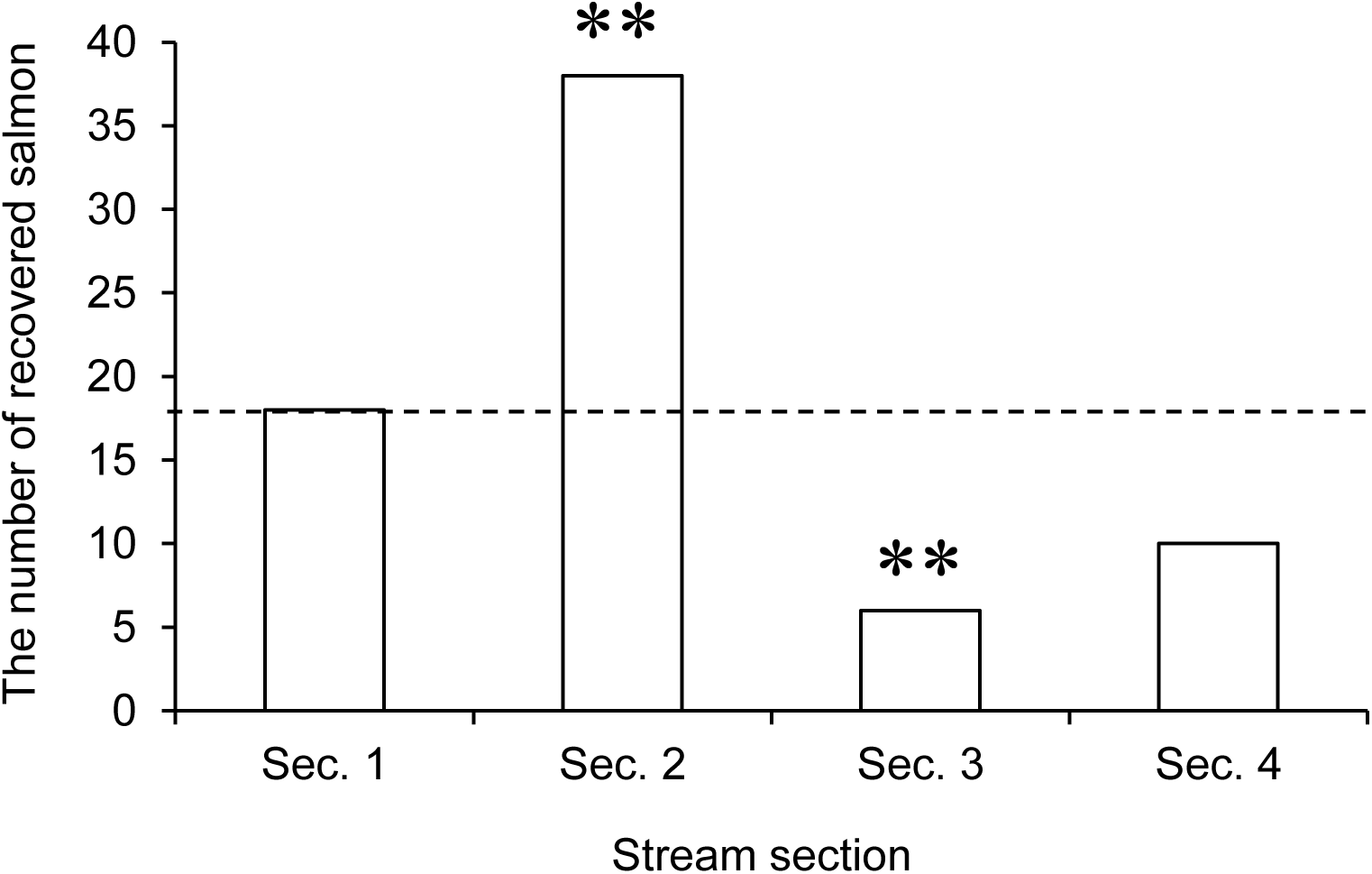
The number of juvenile masu salmon (*Oncorhynchus masou*) recovered at each section in the semi-natural stream. In each section of the stream, fish number was calculated as the sum of fish remaining from each size category (75-100 and 135-160 mm). Broken line indicates the expected surviving proportion (we hypothesized a uniform 1:1:1:1 ratio). Asterisk denotes a significant deviation from the average value (** *p* < 0.010). Each section (Sec.) was named from the upper most section in the stream, Sec. 1, Sec. 2, Sec. 3 and Sec. 4, Sec. 2 was the stocking location.

Cameras captured grey heron, brown dipper (*Cinclus pallasii*), Japanese red fox (*Vulpes vulpes japonica*) and large-billed crow during the predation test in the semi-natural stream. The total number of photos containing potential predators was 470. Among them, 455 contained grey heron. There were fewer than 10 photos of other animals (brown dipper: one times, Japanese red fox: seven times, large-billed crow: seven times). The most grey heron photos (*n* = 104) were taken at Sec. 2 on day-5 (Fig. 5). Of all the pictures that contained grey herons, 76.5 % were taken at Sec.2. Five photos had two grey herons in each picture, and two photos had three grey herons in each picture, all photos with multiple herons were taken in Sec. 2 between day-4 and day-7 after stocking. Seventeen photos showed grey herons capturing prey fish. Grey herons were most frequently photographed in the mornings and evenings.

**Fig. 5.**
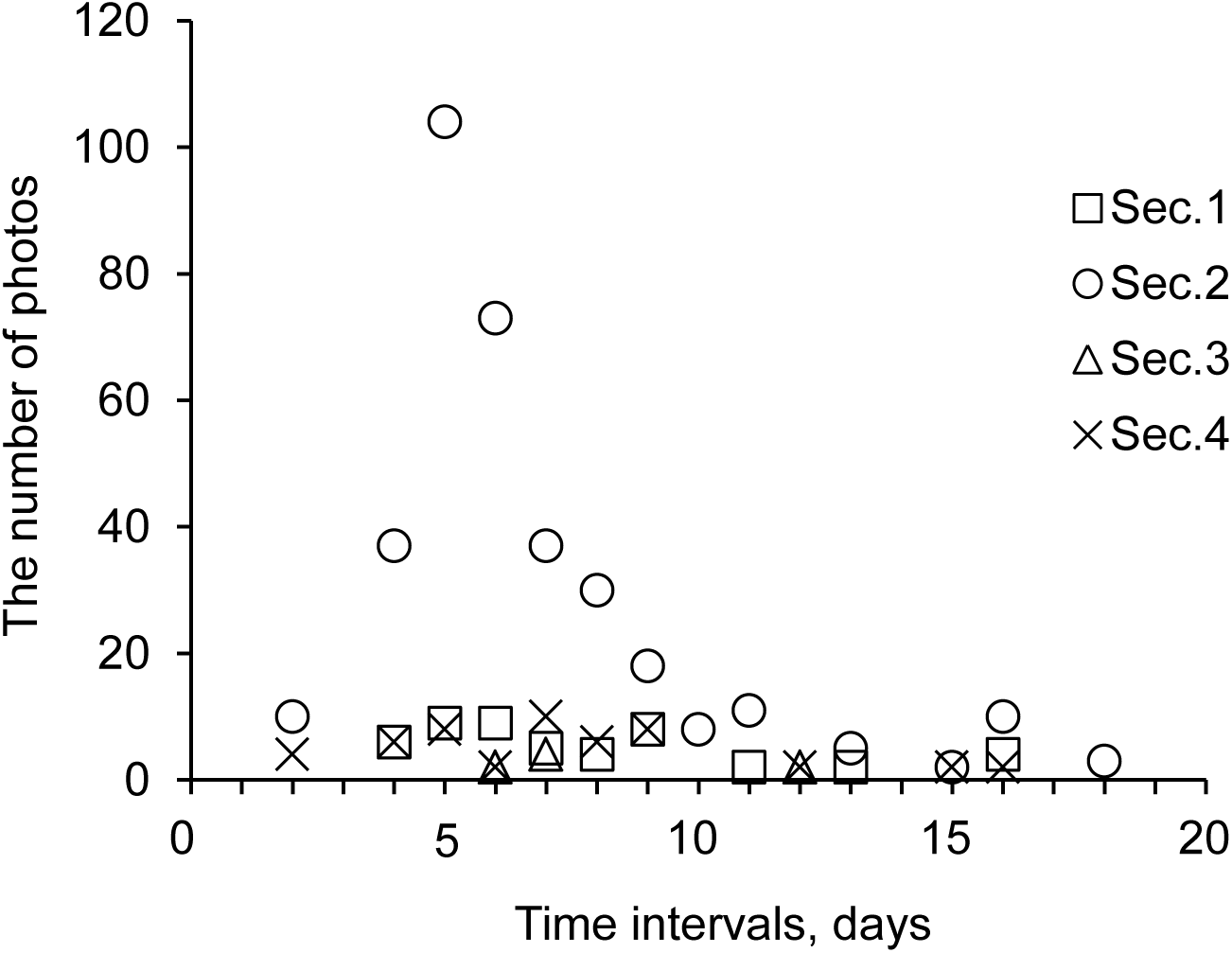
Number of photos of grey heron (*Ardea cinerea*) in each study section of semi-natural stream. X-axis show the number of days elapsed from stocking (day 0) to completion at day 20.

In order to investigate the position that grey herons utilized, 412 photos were analyzed (455 minus 43 photos for undistinguishable positioning). The average proportion of photos showing grey heron around the pools (6.88 ± 10.68 %) (Fig. 1c) was significantly lower than those around the riffles (93.13 ± 10.68 %) (Fig. 1d) (*F*_1, 6_ = 53.13, *P* < 0.010) (Fig. 6). The overall proportions of photos taken of grey herons in the morning (3:00-9:00) and in the evening (15:00-21:00) were 49.3% (*n* = 224) and 47.9% (*n* = 218), respectively.

**Fig. 6.**
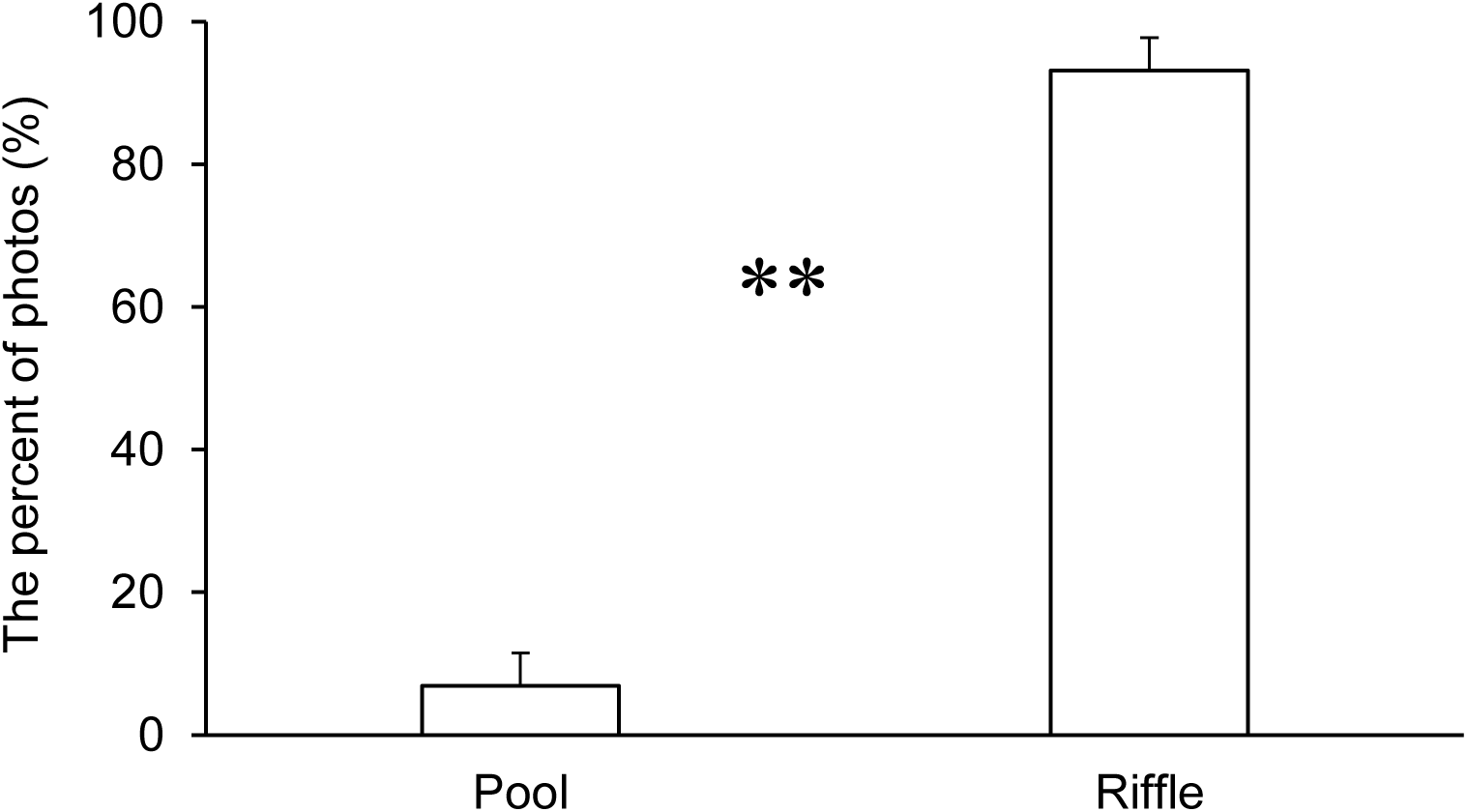
The percent of the photos that showed grey heron located around the pool or riffle across each study section of the semi-natural stream. Asterisk denotes a significant difference (* *p* < 0.050, ** *p* < 0.010). Data are means ± standard error.

## Discussion

In the 20-day stream experiment, 82% of fish were lost and that grey heron was the most frequently visiting predator. It has been previously reported that salmonid populations can be seriously damaged by avian predation (Feltham 1995; Stewart *et al*.2005; Evans *et al*. 2016), but quantitative assessments of the impact of grey heron on river fisheries is rare (Hodgens *et al*. 2004). Our results suggest that at least in our experimental settings, grey herons can significantly reduce salmonid populations by consuming juvenile fish. It is important to note that water depth can strongly influence wading birds’ selection of foraging habitat (Master *et al*. 2005, Gawlik and Crozier 2007). The length of a grey heron’s legs restricts the maximum depth at which they can forage up to 16 cm deep (Ntiamoa-Baidu *et al*. 1998), thereby limiting the habitat suitable for hunting. Our observation that grey herons showed a significant preference for the riffle, rather than the pool, in the semi-natural stream is consistent with previous studies. Thus, where local salmonids reside in shallow water, it is reasonable to expect that grey herons exert a high predation pressure.

On the first day of the tank experiments, grey herons preyed upon large salmon more often than upon small salmon. In both the tank and the stream experiments, the survival rate of large salmon was less than that of small salmon. Birds of the heron family (Ardeidae) including the grey heron are reported to show a preference for larger mosquito fish *Gambusia affinis* (Britton and Moser 1982) and sailfin molly *Poecilia latipinna* (Trexler *et al*. 1994) as prey. It is suggested that larger fish are easier for avian predators to detect than small ones (Eriksson 1985, Magnhagen 1988). After the second day of the tank experiment, however, both sizes of salmon showed similar levels of decline on their survival rate. One possible explanation for this result is the decreased opportunities to prey upon larger fish. Our results suggest that size-selective predation by grey herons depends on the density of preferred prey size, and it appears that size-selective predation occurred in the stream experiment as well as in the tank experiments. Therefore, if fish with different body sizes are stocked in rivers, it might be important to consider how size composition will potentially influence the behavior of predators.

It is established that larger fish are safer from fish-on-fish predation than smaller fish (Peterson and Wroblewski 1984, Houde 1987, Miller *et al*. 1988, Miyamoto and Araki 2017). One potential cause for the higher survival rate of larger fish is improved swimming ability that allows them to better avoid predators as they grow (Beamish 1978; Lundvall *et al*. 1999). However, the grey heron is an ambush predator that usually stands upright and waits for a fish to approach (Tojo, 1996), so the swimming ability that a fish possesses to avoid aquatic predators has significantly limited benefits to them in this case (Miyamoto 2016b), and large size might make fish more detectable. Therefore, to decrease predation risk by wild animals, we suggest that hatchery managers choose an optimum body size of released fish based on the predator species inhabiting the stocking area. Additionally, to block the vision of grey herons or prevent access, it can be recommended to stock fish in a structurally complex place that has a lot of woody debris and/or aquatic vegetation (Diehl 1988) if many grey herons are around during the stocking season.

Generally, in salmonids, larger individuals avoid exposure to a predatory threat and reduce growth rate, whereas smaller individuals are less cautious and maintain their growth rate even in the presence of a threat (Reinhardt 1999). Based on this behavior, it is expected that the OYO salmon are more difficult for grey heron to catch than the YOY salmon. In contrast to this, it was also reported that the hatchery environment selects for bolder individuals that spend more time in open areas and are more active than wild specimens (Sundström *et al*. 2004). So, OYO salmon that spent a longer time than the YOY salmon in the hatchery may display a higher tendency for bold behavior (Roberts *et al*. 2014). Thus, to fully understand the predation risks for stocked salmon, further studies that estimate the individual behavior of fish (e.g. bold and shy) and improve the maladapted behavior of fish (Berejikian *et al*. 1999, Roberts *et al*. 2011) will be needed.

In the tank experiment where different numbers of salmon were stocked, the number of grey herons present or the duration of their sojourn at the tanks was positively correlated to the number of stocked fish. In other studies, large numbers of fish have been observed to remain close to where they were stocked (Cresswell 1981), and piscivorous water birds have displayed similar density and temporal trends in response to stocking (Draulans 1987; Gawlik 2002). This suggests that for the duration of our stream experiment many salmon stayed in Sec. 2, as multiple grey herons simultaneously appeared frequently in that section. We also captured more fish in Sec. 2 by electrofishing than in any other sections on the final day of the stream experiment. These results indicate that most stocked fish gathered without migrating from the initial stocking site and continued to be preyed upon there. This situation may have serious implications for the conservation and propagation of salmonids and it is important to further investigate the spatial and temporal relationships between salmonids and their predators.

The results of this study showed that the grey heron has a tendency to avoid deep water and foraging in the middle of day and night. In contrast, it previously was reported that grey herons forage constantly during the daytime (Richner 1986; Sawara *et al*. 1990), so human activities (Klein, 1993) and other feeding sites (Richner 1986) in the surrounding environment could have had an effect on the observed behavior. Grey herons infrequently forage at night, and it has been suggested that such feeding behavior is inefficient (Sawara *et al*. 1990). Therefore, to decrease the predation pressure of grey herons, just after the fish were released, it might be wise to stock fish in a deep area or at night (Roberts *et al*. 2009), or both. In this way, if we can develop techniques to mitigate predatory damage to fisheries by investigating the relationship between stocked fish and predatory animals, it will not only help the conservation and propagation of stocked fish, but also support the conservation of wild animals as well.

## Acknowledgements

We would like to thank Kouji Mutou, Hidefumi Nakamura and Masaharu Murakami of the National Research Institute of Fisheries Science for their help with the care of the fish. We thank Dr Tsutomu Takeda of Nikko Yumoto Visitor Center of Nikko National Park for his help with the identification of wildlife. We are grateful to Predrag Davidovic and Petar Ilic for critically reading the manuscript and providing valuable comments. Dr Tomoyuki Nakamura of the NRIFS helped us with valuable advice. This work was supported by JSPS KAKENHI Grant Number JP26292102.

